# Tracking changes in corticospinal excitability during visuomotor paired associative stimulation to predict motor resonance rewriting

**DOI:** 10.1101/2025.02.17.638676

**Authors:** Giacomo Guidali, Nadia Bolognini

## Abstract

Mirror properties of the action observation network (AON) can be modulated through Hebbian-like associative plasticity using paired associative stimulation (PAS). We recently introduced a visuomotor protocol (mirror-PAS, m-PAS), which pairs transcranial magnetic stimulation (TMS) over the primary motor cortex (M1) with visual stimuli of ipsilateral (to TMS) movements, leading to atypical corticospinal excitability (CSE) facilitation (i.e., motor resonance) during PAS-conditioned action observation. While m-PAS aftereffects are robust, little is known about markers of associative plasticity during its administration and their predictive value for subsequent motor resonance rewriting.

In the present study, we analyzed CSE dynamics in 81 healthy participants undergoing the m-PAS before and after passively observing left- or right-hand index finger movements. Here, typical and PAS-conditioned motor resonance was assessed with TMS over the right M1. We examined CSE changes during the m-PAS and used linear regression models to explore their relationship with motor resonance modulations.

Results showed that the m-PAS transiently reshaped both typical and PAS-conditioned motor resonance. Importantly, we found a gradual increase of CSE during m-PAS, which predicted the loss of typical motor resonance but not the emergence of atypical responses after the protocol’s administration.

Our findings suggest that the motor resonance reshaping induced by the m-PAS is not fully predictable by CSE online modulations. Likely, this rewriting is the product of a large-scale reorganization of the AON rather than a phenomenon restricted to the PAS-stimulated motor cortex. This study underlines that monitoring CSE during non-invasive brain stimulation protocols could provide valuable insight into some, but not all, their plastic outcomes.

## 1. INTRODUCTION

Associative sensorimotor learning plays a crucial role in shaping mirror properties of the human brain (Cook et al., 2014; Heyes & Catmur, 2022; Keysers & Gazzola, 2014). Recent studies highlight the feasibility of modulating mirror neuron responses using experimental paradigms leveraging this form of learning (e.g., Bardi et al., 2015; Brunsdon et al., 2020; Catmur et al., 2007; Catmur & Heyes, 2019; de Klerk et al., 2015; Fitzgibbon et al., 2016) or the induction of Hebbian-like associative plasticity through non-invasive brain stimulation protocols (Chiappini et al., 2024; Guidali et al., 2020, 2025; Guidali, Picardi, Gramegna, et al., 2023; Maddaluno et al., 2020; Turrini et al., 2024; Zazio et al., 2019).

Considering the domain of action observation, visuomotor properties of human mirror neurons can be studied non-invasively by exploiting the motor resonance phenomenon, i.e., the corticospinal excitability (CSE) enhancement detectable during the observation of biological movements (Craighero, 2024; Fadiga et al., 1995; Naish et al., 2014). This phenomenon is assessed by recording motor-evoked potentials (MEPs) from transcranial magnetic stimulation (TMS) of the primary motor cortex (M1). It is thought to reflect the activation of mirror neuron populations located in the ventral premotor cortex (PMv), a key hub of the *action observation network* (AON - Rizzolatti et al., 2014), which have direct connections with M1, which in turn influence motor system excitability during action observation (e.g., Avenanti et al., 2007; Cantarero et al., 2011; Catmur et al., 2011; Chiappini et al., 2024; de Beukelaar et al., 2016; Koch et al., 2010; Lago et al., 2010).

Motor resonance responses can be experimentally modulated with *paired associative stimulation* (PAS), a class of non-invasive brain stimulation protocols repeatedly coupling a peripheral and a cortical stimulation activating the same cortical area/circuit, in turn inducing Hebbian associative plasticity in the target system (for reviews, see: Guidali et al., 2021a, 2021b; Suppa et al., 2017). Our research group introduced a visuomotor version of the PAS targeting the AON, called the mirror PAS (m-PAS - Guidali et al., 2020), which reliably induces a reshaping of motor resonance responses for simple movements by leveraging their hemispheric lateralization (i.e., MEP increase can be detected only when recorded by stimulating the M1 contralateral to the observed movement – e.g., Aziz-Zadeh et al., 2002). During the m-PAS, TMS pulses over M1 are repeatedly coupled with the observation of visual stimuli depicting a finger movement made with the ipsilateral (to the TMS site) hand, a condition that at baseline is unrelated to CSE facilitation. After m-PAS administration, the successful induction of atypical motor resonance emerges, as indexed by CSE facilitation in the ipsilateral hemisphere when the PAS-conditioned movement is observed (Guidali et al., 2020). Interestingly, this emergence occurred at the cost of the typical response, significantly reduced after the protocol’s administration (Guidali et al., 2025). The m-PAS corticospinal effects are also accompanied by modulations of AON activation behavioral markers (i.e., automatic imitation; Guidali, Picardi, Gramegna, et al., 2023) and TMS-evoked M1 functional connectivity in the alpha and beta bands during action observation (Guidali et al., 2025).

Recently, a cortico-cortical PAS targeting PMv-to-M1 connectivity was also found effective in modulating (typical) motor resonance and automatic imitation (Chiappini et al., 2024; Turrini et al., 2024). As said before, CSE facilitation during action observation is thought to reflect excitatory connections between ventral premotor regions of the AON and M1 (for reviews, see: Fadiga et al., 2005; Rizzolatti et al., 2014), and this explains why targeting such a cortico-cortical connection could influence AON activation proxies (Chiappini et al., 2024; Turrini et al., 2024). In this vein, we can speculate that also the m-PAS recruits a premotor-to-motor pathway, even if not directly stimulated with TMS (Guidali et al., 2020, 2025). Notably, using the PMv-M1 cortico-cortical PAS, Turrini et al. (2022, 2023, 2024) found a gradual enhancement of MEP amplitude during its administration, suggesting that, at least for this ccPAS version, CSE could be used as a reliable, online marker of Hebbian associative plasticity induction within M1. Still, this enhancement was not investigated in relation to the aftereffects of the protocol (i.e., whether CSE modulations during PAS are somehow predictive of the protocol’s outcomes at the single-subject level) (Turrini et al., 2022, 2023, 2024). Hence, assuming a shared neurophysiological substrate between the PMv-M1 cortico-cortical PAS and the m-PAS, CSE could also be modulated similarly during the latter visuomotor protocol. This could reflect the (gradual) induction of associative plasticity within the motor system and, likely, a plastic rewiring of the stimulated M1 inter-areal communication, responsible for the (atypical) AON recruitment at the sight of the PAS-conditioned movement after the protocol’s administration (Guidali et al., 2025). Critically, if CSE modulations during the m-PAS reflect the induction of state-dependent associative plasticity within the AON, we could further hypothesize that their magnitude may predict the reshaping of motor resonance patterns, in turn providing valuable information on the neurophysiological underpinnings of m-PAS plastic modulations and, in a broader perspective, on how AON-driven associative plasticity built up during a visuomotor protocol based on (passive) action observation.

Given these premises, in the present work, we investigate possible CSE changes during m-PAS administration and whether they predict the protocol’s aftereffects on typical and experimentally induced motor resonance. To do this, we have analyzed MEPs recorded online during the m-PAS protocol, aggregating data from previous studies conducted by our research group (Guidali et al., 2020, 2025; Guidali, Picardi, Gramegna, et al., 2023). First, we assessed m-PAS effects in our (aggregated) sample to replicate patterns found for (typical and atypical) motor resonance responses in our previous studies. Then, we analyzed the evolution of the CSE temporal profile during m-PAS administration. Finally, we conducted linear regression models to assess possible relations between motor resonance modulations and changes in CSE during protocol administration. Our investigation aims to shed better light on possible markers of sensorimotor associative plasticity induction during the administration of the m-PAS, exploring whether they could predict the magnitude of subsequent aftereffects.

## 2. MATERIALS and METHODS

### 2.1. Participants

We took datasets of the present work from 93 subjects who participated in a series of previous m-PAS studies conducted by our research group (Guidali et al., 2020, 2025; Guidali, Picardi, Gramegna, et al., 2023). None of these studies had explored possible modulation of CSE during the m-PAS protocol. For the present work, we considered only data from sessions where the m-PAS administered to the participant had the parameters found effective in all our studies (i.e., with an inter-stimulus interval – ISI – between paired stimulations of 25 ms, depicting right-hand index finger movements, and with TMS delivered over right M1). We excluded from our initial dataset participants whose MEPs were not recorded during the m-PAS protocol due to technical issues (n=8). Four participants took part in more than one experiment; hence, we considered data only from the first experiment in which they participated. All this considered, the final sample included 81 right-handed healthy participants (35 males, mean age ± standard deviation – SD: 23.7 ± 2.9 years; mean education ± SD: 15.6 ± 1.9 years; mean Edinburgh handedness inventory score ± SD: 74 ± 15.9%). To assess whether this number of participants is sufficient to obtain reliable results from linear regression analyses, we run an a-priori power analysis with the software G*Power 3.1 (Faul et al., 2009). The power analysis [*f ^2^* = .15 – corresponding to a medium desired effect size (Cohen, 2013), alpha error level: *p* = .05; statistical power = .9, actual power = .9] suggested at least 73 participants to achieve enough statistical power. All the original experiments were performed following the ethical standards of the Declaration of Helsinki and after the approval of the Ethical Committee of the University of Milano-Bicocca. Before taking part in the study, participants gave their written informed consent. Dataset and analyses of the present study will be publicly available on Open Science Framework (OSF) once the manuscript will be accepted for publication.

### 2.2. m-PAS and action observation task

The m-PAS is a visuomotor version of the PAS protocol where a TMS pulse over the right M1 is repeatedly coupled with a moving right hand (hence ipsilateral to stimulation) to promote the induction of atypical motor resonance responses (Guidali et al., 2020, 2021b). The m-PAS consisted of 180 paired stimulations delivered at a rate of 0.2 Hz (total duration: 15 min). Each trial started with a frame depicting a right hand in an egocentric perspective at rest. After 4250 ms, a second frame depicting the same hand performing an abduction movement with the index finger appeared for 750 ms, giving rise to apparent motion. After 25 ms from the onset of this frame, we delivered a TMS pulse over the right M1 at 120% individual resting motor threshold (rMT – **Figure 1a**). The correct timing of the frames was checked by using a photodiode.

**Figure 1.**
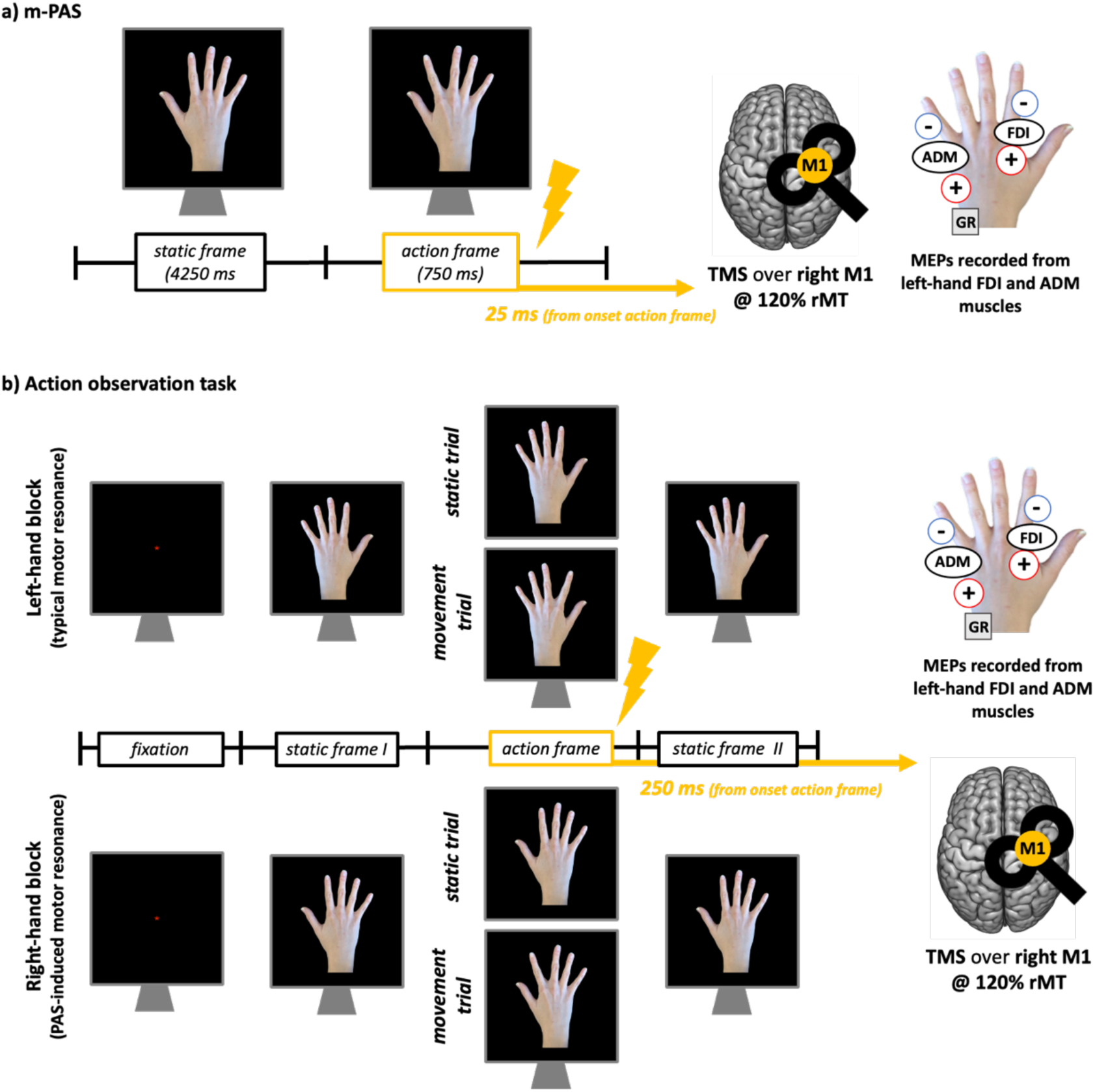
m-PAS protocol (**a**) and action observation task (**b**) used in the datasets considered for the present work. In the action observation tasks, the timing of the frames slightly varied according to the study from where MEP data were taken (see: Guidali et al., 2020; 2023, 2025 for further details). M-PAS parameters were the same in every study.

In all our datasets, we assessed motor resonance before and after m-PAS administration during a standard passive action observation task where participants had to observe, in separate blocks, a left or a right hand performing index finger abduction movements (Guidali et al., 2020, 2025; Guidali, Picardi, Gramegna, et al., 2023). As for the m-PAS, the rapid succession of two frames – one depicting the hand at rest (static frame) and the other depicting it performing the movement (action frame) – gave the illusion of apparent motion. Regardless of the block (i.e., depicting left or right hands), TMS was always delivered over the right M1 at 120% rMT. During static trials, TMS was delivered while participants observed the hand at rest. During movement trials, TMS was delivered 250 ms from the onset of the action frame, depicting the abduction movements of the index finger (**Figure 1b**). Half of the trials in each block depicted static hands and half moving ones. We instructed participants to keep their hands relaxed and out of view and carefully observe these visual stimuli and, as an attentive task similar to control that participants were looking at the PC screen, to respond when a red dot appeared on the depicted hand (10% of total task trial, which we excluded from following analysis). Trial duration and the total number of stimuli in each block slightly varied across studies, and we referred the reader to the original works for more details (Guidali et al., 2020, 2025; Guidali, Picardi, Gramegna, et al., 2023); each block lasted about 5 min.

M-PAS and action observation tasks were under computer control, running on the E-Prime software (E-Prime 2.0, Psychology Software Tool, Inc.).

### 2.3. TMS

We delivered TMS pulses with a biphasic figure-of-eight coil (diameter = 70 mm) connected to a Magstim Rapid 2 (Magstim, Whitland, UK) or a Nexstim Eximia stimulator (Nexstim, Helsinki, Finland). At the beginning of the session, we found the motor hotspot of the left FDI muscle by moving the coil in 5 mm steps around the presumed right-hemisphere motor hand area by using a slightly supra-threshold stimulus and recording MEPs. Based on the specific experiment, the individual rMT was determined either as the lowest TMS intensity (expressed as a percentage of the maximum stimulator output) that could evoke an MEP of at least 50 µV in the FDI muscle in 5 out of 10 trials (Rossi et al., 2009) or using the parameter estimation by sequential testing (PEST) method (Awiszus, 2003). On average, the mean rMT of our sample was (mean ± SD) 44.4 ± 10.8% (with no statistically significant difference concerning the procedure adopted, *t* = .231, *p* > .673). The stable TMS coil placement and position were constantly monitored during the experimental sessions through neuronavigation software, i.e., SofTaxic Optic 2 (EMS, Bologna, Italy) for data collected with the Magstim Rapid 2 stimulator (Guidali et al., 2020), and the integrated navigated brain stimulation system (Nexstim, Helsinki, Finland) for data collected with the Nexstim Eximia stimulator (Guidali et al., 2025; Guidali, Picardi, Gramegna, et al., 2023). The coil was consistently positioned tangential to the scalp and angled at 45° to the midline, perpendicular to the targeted cortical gyrus. This orientation generated brain currents in the anterior-to-posterior (first phase)/posterior-to-anterior (second phase) direction.

### 2.4. Electromyographic (EMG) recording and preprocessing

In all our datasets, we recorded MEPs from FDI (target muscle, implicated in the index finger movements observed by participants) and ADM muscles (control muscle, not implicated in the movements observed) of the left hand with Signal software (version 3.13, Cambridge Electronic Devices, Cambridge, UK). EMG signal was sampled at 5000 Hz using a Digitimer D360 amplifier (Digitmer Ltd, Welwyn Garden City, UK) connected to a CED micro1401 A/D converter (Cambridge Electronic Devices, Cambridge, UK). Active electrodes were placed over the muscle bellies; reference ones over the metacarpophalangeal joint of the index (for FDI) and little finger (ADM). The ground electrode was placed over the right head of the ulna. EMG signal was amplified, band-pass (10–1000 Hz), notch filtered, and stored for offline analysis. Data were collected from 100 ms before to 200 ms after the TMS pulse (time window: 300 ms). We analyzed MEPs offline using Signal software (version 3.13). MEP peak-to-peak amplitude was calculated in each trial of the m-PAS and the action observation task between 5 ms and 60 ms from the TMS pulse. Trials with artifacts deviating from 100 µV in the 100 ms before the TMS pulse and trials in which MEP amplitude was smaller than 50 µV were excluded from the analysis.

### 2.5. Statistical analyses

For action observation tasks, a *motor resonance index* (Guidali et al., 2025; Guidali, Picardi, Franca, et al., 2023) was computed as the ratio in MEP amplitude between movement and rest trials:

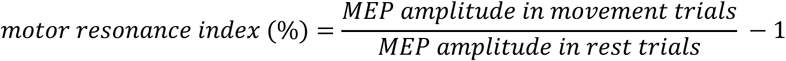

Namely, for every participant and condition, the mean MEP amplitude in trials depicting the movement was divided for MEP amplitude from rest trials of the same condition, which served as a baseline for CSE. The value ‘1’ was subtracted from the ratio to express the percentage concerning the rest condition. Hence, positive values indicated corticospinal excitability facilitation by action observation (and thus the presence of motor resonance). MEP data recorded during the m-PAS were normalized using z-point transformation, divided into 6 bins of 30 trials each (i.e., *bin 1* = trials 1-30; *bin 2* = trials 31-60; *bin 3* = trials 61-90; *bin 4* = trials 91-120; *bin 5* = trials 121-150; and *bin 6* = trials 151-180), and the mean peak-to-peak amplitude in each bin was calculated to explore the temporal evolution of CSE during the protocol. Raw MEP values for each muscle are reported in **Supplemental Tables 1** and **2**, separated for the action observation task’s conditions and m-PAS bins.

First, as a quality check to assess – and replicate – the effect of the m-PAS on motor resonance (Guidali et al., 2020, 2025; Guidali, Picardi, Gramegna, et al., 2023), we run a ‘viewed Hand’ (left-hand, right-hand) X ‘Time’ (pre-PAS, post-PAS) X ‘Muscle’ (FDI, ADM) repeated-measures analysis of variance (rmANOVA) on the *motor resonance index* values. Then, we assessed whether CSE changed during m-PAS administration with a ‘Bin’ (1, 2, 3, 4, 5, 6) X ‘Muscle’ (FDI, ADM) rmANOVA on normalized MEP amplitude. Finally, given the significative patterns found (see **Results**), we performed a series of linear regression analyses for each muscle, separated for left- and right-hand observation conditions, aiming to explore whether changing in CSE during the m-PAS predicted protocol’s aftereffects on typical and atypical motor resonance. Here, we considered the difference between *motor resonance index* values after and before m-PAS administration (i.e., *motor resonance gain*) as the dependent variable and the difference between normalized MEP amplitude in the final and first bin of the m-PAS (i.e., *m-PAS CSE gain*) as the predictor.

All statistical analyses were performed using the software Jamovi (The Jamovi Project, 2025). Statistical significance was set at *p* < .05. Normality of the distributions was confirmed for all our variables, checking it with the Shapiro-Wilk test and Q-Q plots assessment. For rmANOVAs, data sphericity was tested by applying Mauchly’s test in every dataset. When data sphericity was not confirmed, the Greenhouse-Geisser correction was applied. Significant effects were further explored with Tukey HSD-corrected post-hoc comparisons if not otherwise specified. Partial eta-squared (*η_p_^2^*), Cohen’s *d*, and coefficient of determination (*R^2^*) were calculated in every rmANOVA, t-test, and regression, respectively, and reported as effect size values. For linear regressions, also the standardized regression coefficient (*β*) was report. In the **Results** section, mean ± SE is reported for each variable.

## 3. RESULTS

### 3.1. Motor resonance patterns before and after m-PAS administration

Results from the rmANOVA on the *motor resonance index* to check CSE patterns after m-PAS administration showed a significant ‘viewed Hand’ X ‘Time’ X ‘Muscle’ interaction (*F*_1,80_ = 15.57, *p* < .001, *η_p_^2^* = .16), as well as main effect of factor ‘Muscle’ (*F*_1,80_ = 5.23, *p* = .025, *η_p_^2^* = .06) and interaction ‘viewed Hand’ X ‘Time’ (*F*_1,80_ = 21.29, *p* < .001, *η_p_^2^* = .21). No other significant effect was found (all *Fs* < 2.17, all *ps* > .144). Motor resonance patterns were further explored with two separate rmANOVA, one for each muscle, as previously done in all our works (Guidali et al., 2020, 2023, 2024).

For the FDI muscle, this analysis showed only a significant main effect of the ‘viewed Hand’ X ‘Time’ interaction (*F*_1,80_ = 52.93, *p* < .001, *η_p_^2^* = .4). No other significant effect was found (all *F*s < .46, all *p*s > .507). Post-hoc comparisons showed that, as expected, motor resonance at baseline is present only for left-hand conditions (mean *motor resonance index* ± standard error: 12.78 ± 1.73%); right-hand conditions did not give rise to motor resonance (.69 ± 1.36%; *vs.* pre-PAS left-hand *motor resonance index*: *t*_80_ = 5.57; *p* < .001, *d* = .62). After the administration of the m-PAS, atypical motor resonance for the PAS-conditioned right-hand movements emerged (12.83% ± 1.48%; *vs.* pre-PAS right-hand *motor resonance index*: *t*_24_ = 6.63, *p* < .001, *d* = .74) accompanied with a rewriting of the typical phenomenon. Indeed, after the m-PAS, the *motor resonance index* for left-hand conditions (4.83 ± 1.16%) was significantly lower than in baseline (*t*_24_ = -3.86, *p* = .001, *d* = -.43) and from the one obtained for right-hand stimuli (*t*_24_ = -4.57, *p* < .001, *d* = -.51; **Figure 2a**).

**Figure 2.**
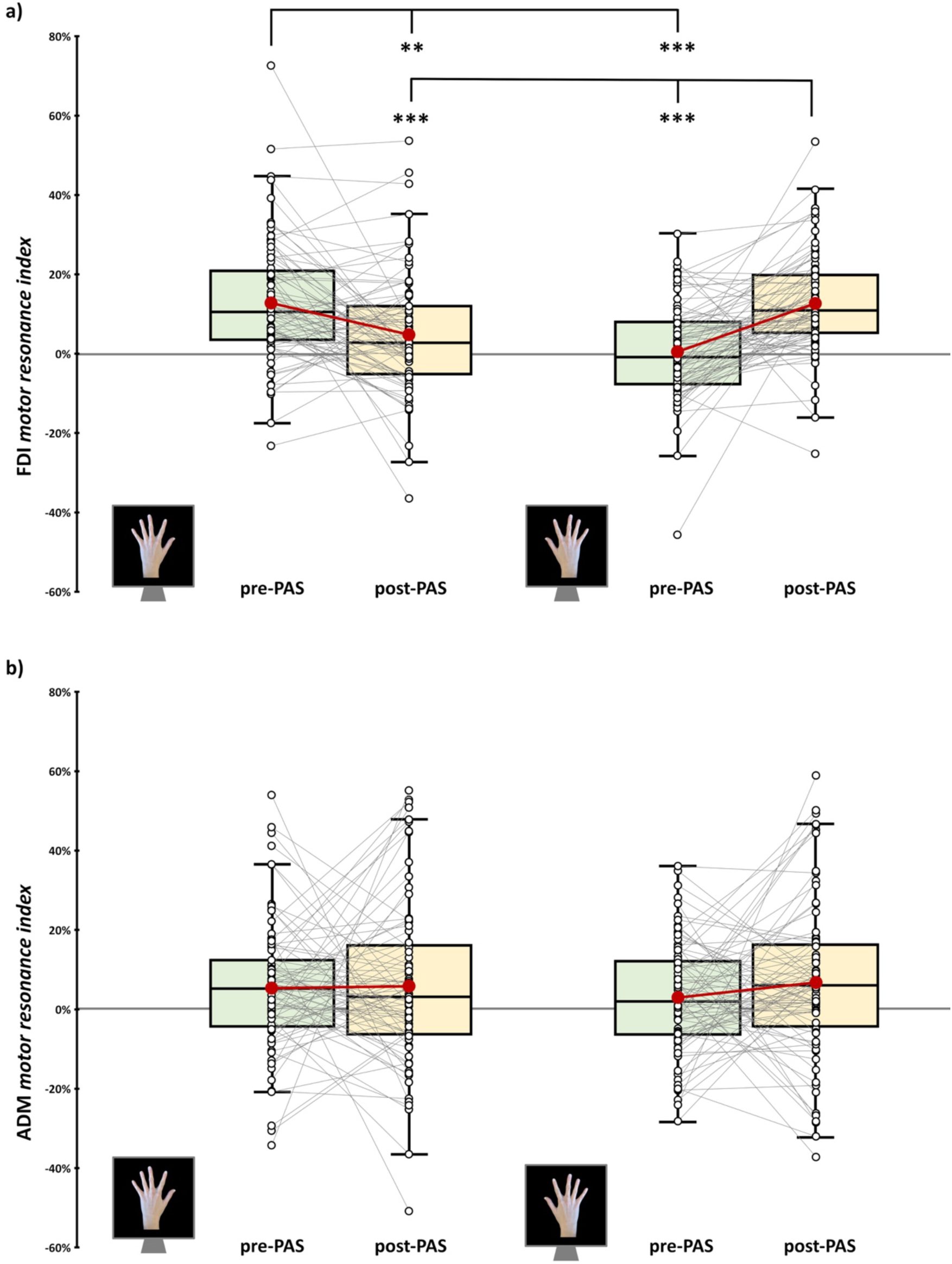
*Motor resonance index* before (green boxplot) and after (yellow boxplot) m-PAS administration for FDI (**a**) and ADM (**b**) muscles. In the box-and-whiskers plots, red dots indicate the means of the distributions. The center line denotes their median values. Black-and-white dots show individual participants’ scores. The box contains the 25^th^ to 75^th^ percentiles of the dataset. Whiskers extend to the largest observation falling within the 1.5 * inter-quartile range from the first/third quartile. Significant *p* values of Tukey corrected post-hoc comparisons are reported (** = *p* < .01; *** = *p* < .001).

For the ADM muscle – not involved in the observed index finger movement, hence acting as a control for the muscle-specificity of motor resonance patterns (Craighero, 2024; Guidali et al., 2020) – the rmANOVA showed no significant effects of factors ‘viewed Hand’ (*F*_1,80_ = .14, *p* = .714, *η_p_^2^* < .01), ‘Time’ (*F*_1,80_ = 1.03, *p* = .314, *η_p_^2^* = .01) and their interaction (*F*_1,80_ = .78, *p* = .38, *η_p_^2^* = .01) (**Figure 2b**).

### 3.2. CSE during m-PAS protocol

rmANOVA on (z-transformed) MEPs recorded during the m-PAS showed a significant effect of main factor ‘Bin’ (*F*_3.1,255.3_ = 13.48, *p* < .001, *η_p_^2^* = .14) but not of ‘Muscle’ (*F*_1,80_ = 2.83, *p* = .096, *η_p_^2^* = .03) or ‘Bin’ X ‘Muscle’ interaction (*F*_3.5,279.6_ = 1.78, *p* = .142, *η_p_^2^* = .02). Namely, CSE changed during m-PAS administration but with patterns that are not muscle-specific. Post-hoc comparisons showed that MEPs recorded during the last 30 trials of the m-PAS protocol (i.e., *Bin 6*) were significantly higher for all the other bins (*vs. Bin 1*: *t*_80_ = 5.36; *p* < .001, *d* = .6; *vs. Bin 2*: *t*_80_ = 5.08; *p* < .001, *d* = .56; *vs. Bin 3*: *t*_80_ = 3.86; *p* = .003, *d* = .43; *vs. Bin 4*: *t*_80_ = 4.74; *p* < .001, *d* = .53; *vs. Bin 5*: *t*_80_ = 3.01; *p* = .039, *d* = .34). Furthermore, MEPs in *Bin 5* significantly differed from ones in the first (*t*_80_ = 3.97; *p* = .002, *d* = .44) and second bin (*t*_80_ = 3.53; *p* = .009, *d* = .39). Overall, this pattern suggested a progressive increase of CSE during the m-PAS for both FDI and ADM muscles (**Figure 3**).

**Figure 3.**
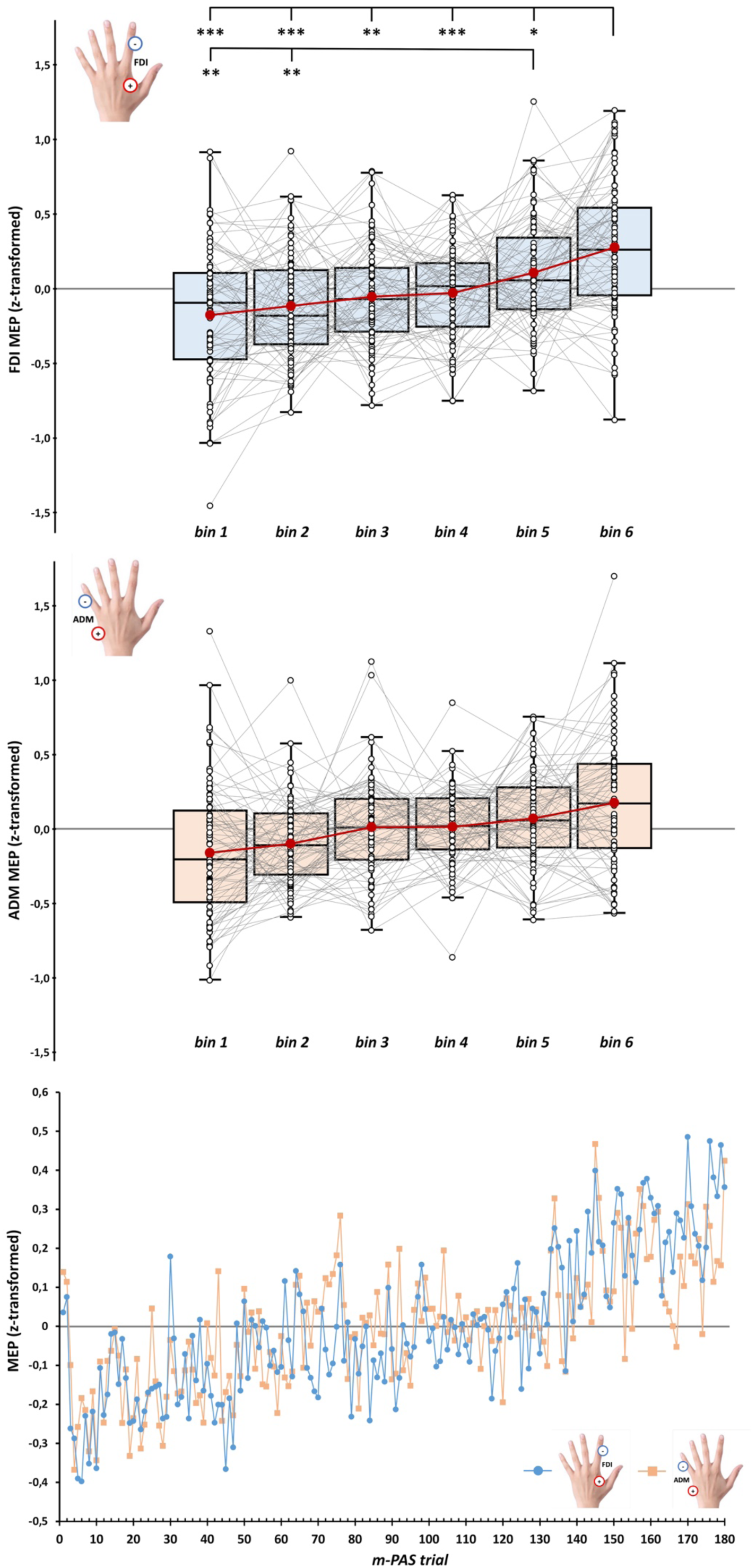
CSE temporal profile during the m-PAS for FDI (upper panel) and ADM (middle panel) muscles according to the 6 bins of 30 trials in which we divided the 180 trials of the m-PAS protocol. Lower panel: mean MEP amplitude at the single-trial level for FDI (blue circles) and ADM (light orange squares). In the box-and-whiskers plots, red dots indicate the means of the distributions. The center line denotes their median values. Black-and-white dots show individual participants’ scores. The box contains the 25^th^ to 75^th^ percentiles of the dataset. Whiskers extend to the largest observation falling within the 1.5 * inter-quartile range from the first/third quartile. Significant *p* values of Tukey corrected post-hoc comparisons for the main effect ‘Bin’ are reported (* = *p* < .05; ** = *p* < .01; *** = *p* < .001).

### 3.3. CSE changes during m-PAS and their relation with motor resonance modulations

Linear regressions run to explore whether changes in FDI CSE during the m-PAS (*m-PAS CSE gain,* i.e., mean normalized MEP amplitude in the first bin subtracted to the one in the last bin) predicted the *motor resonance gain* during the observation of left- or right-hand movements (i.e., *motor resonance index* before the m-PAS subtracted to values found after its administration) showed a statistically significant relation only for left-hand motor resonance (*β* = -.33; *F*_1,79_ = 9.84; *p* = .002; *R^2^* = .11) but not for right-hand one (*β* = .11; *F*_1,79_ = 1.01; *p* = .318; *R^2^* = .01; **Figure 4a**). Namely, the difference between the first and last MEP bin recorded during the m-PAS significantly predicted the loss of typical motor resonance (*motor resonance gain* = -.08 + -.087 * *m-PAS CSE gain*). At variance, the emergence of atypical motor resonance is not associated with CSE modulation during the m-PAS.

**Figure 4.**
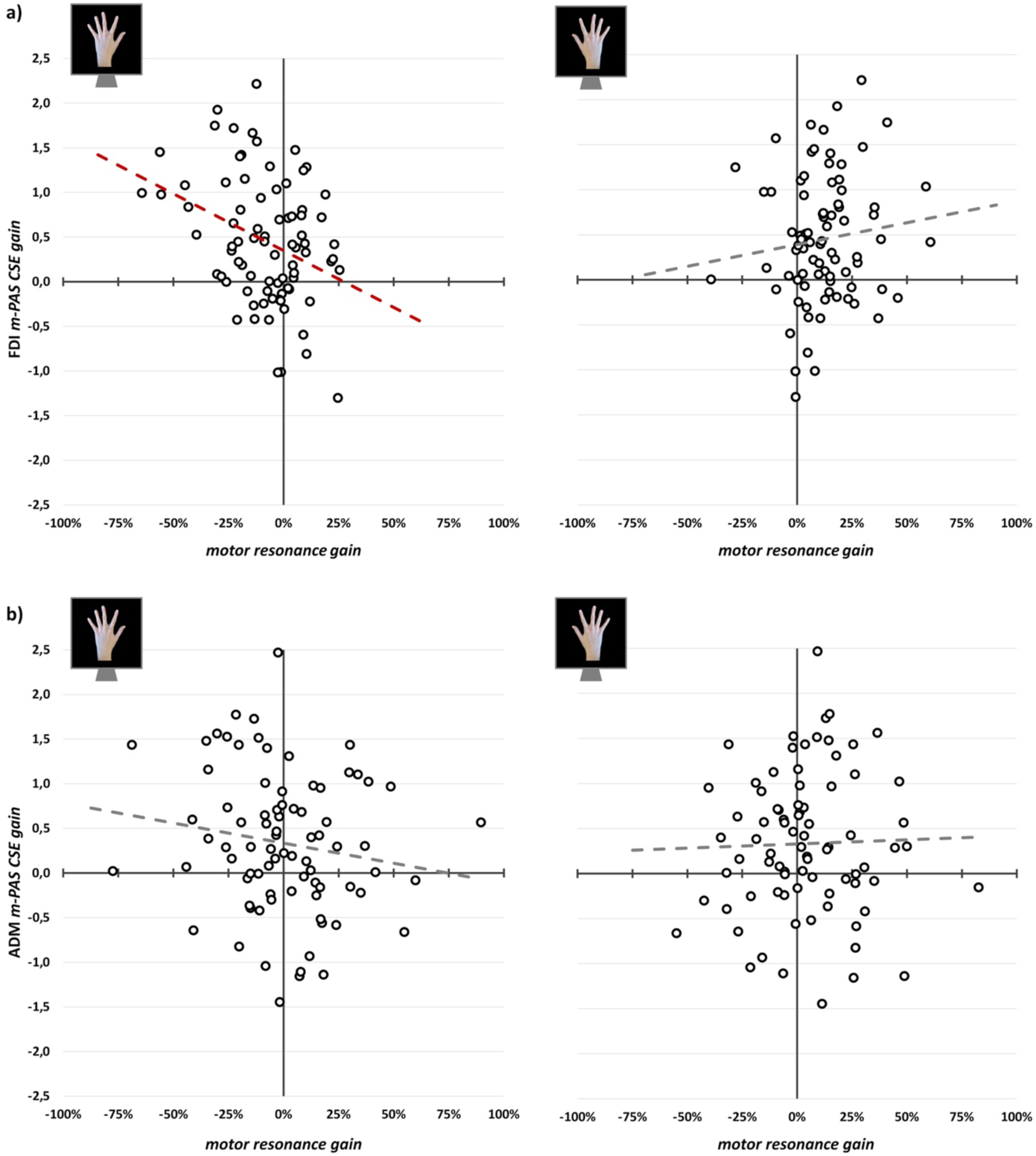
Scatterplots between *m-PAS CSE gain* (i.e., mean normalized MEP amplitude in m-PAS bin 1 subtracted to the one in bin 6) and *motor resonance gain* (i.e., *motor resonance index* before the m-PAS subtracted to values found after its administration) for FDI (**a**) and ADM (**b**) muscles. Dashed lines indicate the linear regression’s fitted line (in red, the significant one found between *m-PAS CSE gain* and left-hand *motor resonance gain* for FDI MEPs).

Given the significant modulation of CSE during m-PAS found for this muscle, we run the same regressions for ADM gain indexes. For this muscle, we found that the *m-PAS CSE gain* did not predict motor resonance gain either during the observation of left- (*β* = -.15; *F*_1,79_ = 1.93; *p* = .169; *R^2^* = .02) or right-hand index finger movements (*β* = .03; *F*_1,79_ = .06; *p* = .803; *R^2^* < .01; **Figure 4b**), highlighting the muscle-specificity of the relation previously found.

## 4. DISCUSSION

Our study investigates whether changes in CSE during a visuomotor version of the PAS (the m-PAS; Guidali et al., 2020) predict the protocol’s aftereffects on typical and PAS-induced motor resonance. We found a gradual enhancement of CSE during m-PAS administration, whose magnitude predicts modulation patterns on typical motor resonance (i.e., loss of CSE facilitation during contralateral movement observation) but not on the atypical, PAS-conditioned phenomenon (i.e., emergence of CSE facilitation during the observation of ipsilateral movements). Overall, these results provide valuable information on how Hebbian associative plasticity within the AON is built up during the protocol, shedding better light on the possible neurophysiological substrates grounding the m-PAS effectiveness.

### 4.1. Motor resonance responses are reshaped after the m-PAS

By aggregating data taken from our previous works with the m-PAS (Guidali et al., 2020, 2025; Guidali, Picardi, Gramegna, et al., 2023), the first novel result is that m-PAS induces not only the emergence of atypical, muscle-specific motor resonance for the movement conditioned during the protocol, but it also disrupts the typical visuomotor association, reducing the magnitude of motor resonance brought about by observing movements performed with the contralateral hand. That is, the m-PAS not only drives the emergence of motor resonance for action performed with limbs ipsilateral to the stimulated motor cortex, but it also concurrently inhibits motor resonance for action performed with contralateral limbs. This pattern was solely documented in our more recent work (Guidali et al., 2025), while, in our previous studies (Guidali et al., 2020, 2023), only trends for this double aftereffect were seen, likely due to the need for a greater sample size to achieve statistical significance.

As already argued (Guidali et al., 2025), acquiring novel motor and visuomotor responses can occur at the cost of the consolidated one (e.g., Catmur et al., 2007, 2011; Cavallo et al., 2014; Hamel et al., 2024; Kobayashi et al., 2009; Takeuchi et al., 2012). Similarly, after m-PAS administration, a novel motor resonance response emerged while observing the conditioned movement, but this atypical reshaping led to a transient disruption of visuomotor responses already in place in the AON (Guidali et al., 2025). This evidence points out that the m-PAS, through associative plasticity induction within the AON, transiently induces a complex reorganization of hemispheric-lateralized motor resonance for simple movements, extending beyond the visual stimulus conditioned during the protocol.

### 4.2. Gradual online enhancement of CSE during the m-PAS

The second key finding is that CSE is significantly modulated during the m-PAS. Notably, this enhancement is detectable from both FDI and ADM muscles, indicating a general increase of motor cortex excitability during m-PAS. This cortical excitability enhancement occurs specifically during the protocol’s administration. In fact, the analysis of raw MEP amplitude collected during the action observation task did not show any modulation of CSE after m-PAS. Still, m-PAS effects were detected only at the level of motor resonance (see **Supplemental Analysis 1**). This evidence means that the online CSE enhancement induced by m-PAS dissociates from its offline effects on motor resonance.

The increase of CSE during the m-PAS aligns well with previous studies using cortico-cortical PAS targeting PMv-M1 connectivity and showing a gradual enhancement of M1 excitability during this protocol (Turrini et al., 2022, 2023), accompanied by modulation of AON activation proxies after its administration (Chiappini et al., 2024; Turrini et al., 2024). As already said in the **Introduction**, the PMv-M1 pathway is the neurophysiological underpinning binding AON and M1 activations and grounding the motor resonance phenomenon (Rizzolatti et al., 2014). Besides studies with the cortico-cortical PAS, different works using repetitive and paired-pulse TMS showed that the perturbation of PMv impacts MEP facilitation during action observation (e.g., Avenanti et al., 2007; Cantarero et al., 2011; Catmur et al., 2011; de Beukelaar et al., 2016). Hence, it is reasonable to suppose its involvement also during the m-PAS, even if not directly stimulated (Guidali et al., 2020). In this framework, the increased CSE during the protocol could be interpreted as evidence that the PMv-M1 pathway is salient for associative plasticity induction during the m-PAS.

An alternative, yet complementary, hypothesis considers the possible contribution of predictive mechanisms (Keysers & Gazzola, 2014; Kilner et al., 2007) during m-PAS administration. Action observation stimuli presented during the m-PAS are highly predictable, considering that participants observed for 15 minutes the same visual stimulus of movement always presented every 5 seconds. As the protocol progresses, the AON can likely be engaged in advance, even before the actual observation of the depicted actions. This is corroborated by previous AON studies with humans and primates showing action anticipation phenomena driven by the presentation of sufficient contextual cues about the movement to be seen (e.g., Aglioti et al., 2008; Maranesi et al., 2014; Qin et al., 2023; Southgate et al., 2009; Urgesi et al., 2010). Furthermore, similar anticipatory mechanisms have been shown to be recruited during a cross-modal version of the PAS, pairing the sight of touch with somatosensory cortical stimulation (Maddaluno et al., 2020; Zazio et al., 2019). In this view, the gradual enhancement of CSE could underlie, besides the induction of state-dependent associative plasticity within the motor system likely mediated by the PMv-M1 pathway, an anticipatory M1 engagement by action observation with the protocol progression.

This evidence stresses that the m-PAS acts at a network level (Guidali et al., 2025) rather than influencing a single AON pathway (like, e.g., cortico-cortical PAS targeting PMv-M1 connectivity; Hernandez-Pavon et al., 2023). From this perspective, our findings suggest that the gradual enhancement of CSE during m-PAS administration would reflect greater M1 reactivity, with associative plasticity being progressively promoted within the entire AON. If this is true, a dynamic reshaping of AON properties could already occur during the protocol, opening up to the chance that CSE modulations detected online could inform the protocol’s aftereffects.

### 4.3. CSE increase during m-PAS predicts only the rewriting of typical motor resonance

The latter hypothesis is confirmed by the third significant result of our study: the magnitude of CSE increase during the m-PAS predicts muscle-specific aftereffects on typical motor resonance; namely, its reduction when viewing action performed with the contralateral hand. This relation is specific for FDI, i.e., the muscle involved in the observed index-finger movement. Hence, even if CSE patterns during the m-PAS are not muscle-specific, reflecting a gradual enhancement of motor system excitability unrelated to the features of the conditioned visual stimulus, their relation with motor resonance modulations follows the typical muscle-specific feature of this phenomenon (Craighero, 2024; Naish et al., 2014). On the contrary, the emergence of the experimentally-induced novel motor resonance effect is unrelated to the online CSE enhancement.

Altogether, these results (i.e., online CSE change predicts the disruption of ‘in place’ visuomotor associations, but not the building up of new ones) suggest that the rewriting of typical motor resonance already occurs during the m-PAS in the targeted motor system, while the induction of new visuomotor association likely took place later, i.e., once the protocol ended. It follows that the CSE enhancement is a successful predictor of some, but not all, the complex modulations of AON activity induced by the m-PAS, likely reflecting Hebbian associative plasticity induction. In this regard, TMS-induced plasticity could also take place in the unstimulated M1. For instance, different clinical studies showed a functional reorganization of inter-hemispheric M1-M1 communication following rehabilitative training based on action observation treatments or mirror therapies (e.g., Mekbib et al., 2020; Michielsen et al., 2011; Novaes et al., 2018; Rizzolatti et al., 2021; Zhang et al., 2024). Thus, it is reasonable to speculate bilateral recruitment of the AON by the m-PAS, resulting in modulation of the contralateral (here the left) M1 excitability that might explain the protocol’s aftereffects on the experimentally-induced motor resonance.

### 4.4. Conclusions and future directions

Overall, our findings corroborate the evidence that m-PAS effects on motor resonance responses are the product of a network-wide reorganization of AON functioning, impacting a vast range of cortical regions with modulations at the CSE level already trackable during the protocol administration. Crucially, the online CSE enhancement can be used to predict just a part of m-PAS aftereffects; in particular, solely modulations more strictly related to the typical, hemispheric-specific motor cortex recruitment during action observation. This evidence aligns well with the results of our recent work integrating m-PAS and neuroimaging, which showed a complex, brain-wide, and frequency-specific reorganization of right M1 functional connectivity during action observation after m-PAS administration, suggesting that typical and experimentally induced motor resonance are not superimposable phenomena, relying on distinct oscillatory dynamics and connectivity patterns (Guidali et al., 2025).

All this considered, the results of the present work open up to intriguing and thought-provoking future directions. For instance, studies could introduce hybrid, cortico-cortical versions of the m-PAS where single-pulse TMS over M1 is replaced with paired-pulse targeting specific AON nodes to disentangle better the contribution of specific pathways in the effects found on CSE and motor resonance during and after m-PAS. At the same time, the contribution of specific AON nodes and pathways could be further explored by varying the features of the visual stimulus of movement conditioned, e.g., replacing the intransitive, simple movement with more ecological and goal-directed actions, which are known to recruit the AON to a greater extent (Aziz-Zadeh et al., 2006; Molenberghs et al., 2012). Similarly, less foreseeable versions of the m-PAS could be tested, assessing the influence of the protocol’s predictability on online and offline markers of associative plasticity induction.

These findings could then open up the introduction of m-PAS versions in clinical settings, with parameters tailored to the clinical profile of single patients. In this vein, a recent proof-of-principle study on stroke survivors suggests that PAS protocols acting on visuomotor mirroring can induce associative plasticity within a damaged motor system, exploiting AON vicarious recruitment (Picardi et al., 2024).

## SUPPLEMENTARY MATERIALS

**Supplemental Table 1.** MEP amplitude raw data (mean ± standard error) from FDI and ADM muscles in the four trial typologies of the action observation task before and after m-PAS administration.

**Supplemental Table 2.** MEP amplitude raw data (mean ± standard error) from FDI and ADM muscles in the 6 bins in which we divided the 180 trials of the m-PAS.

**Supplemental Analysis 1.** rmANOVA on raw MEP amplitude recorded during the action observation tasks.

## CRedIT AUTHOR CONTRIBUTION

**Giacomo Guidali:** conceptualization, methodology, investigation, formal analysis, software, visualization, data curation, writing – original draft

**Nadia Bolognini:** methodology, supervision, funding acquisition, writing – original draft

## INSTITUTIONAL REVIEW BOARD STATEMENT

The study was conducted according to the guidelines of the Declaration of Helsinki, and all the original studies from which we took our data were approved by the Ethics Committee of the University of Milano-Bicocca.

## INFORMED CONSENT STATEMENT

Informed consent was obtained from all subjects involved in the study.

## DATA AVAILABILITY STATEMENT

Dataset and analyses of the present study will be publicly available on Open Science Framework (OSF) once the manuscript will be accepted for publication.

