## Supplemental files for "Tracking changes in corticospinal excitability during visuomotor paired associative stimulation to predict motor resonance rewriting"

**- SUPPLEMENTARY MATERIALS -**

| **Muscle** | **Trial** | **pre-PAS (mV)** | **post-PAS (mV)** |
| --- | --- | --- | --- |
| **FDI** | left-hand static | 1.63 ± .11 | 1.74 ± .12 |
|  | left-hand movement | 1.81 ± .12 | 1.79 ± .12 |
|  | right-hand static | 1.82 ± .11 | 1.57 ± .09 |
|  | right-hand movement | 1.81 ± .11 | 1.77 ± .1 |
| **ADM** | left-hand static | 1.04 ± .08 | 1.05 ± .1 |
|  | left-hand movement | 1.09 ± .09 | 1.07 ± .09 |
|  | right-hand static | 1.02 ± .08 | 1.04 ± .1 |
|  | right-hand movement | 1.04 ± .1 | 1.09 ± .1 |

**Supplemental Table 1.** MEP amplitude raw data (mean ± standard error) from FDI and ADM muscles in the four trial typologies of the action observation task, before and after m-PAS administration.

| **Muscle** | **Bin 1 (mV)** | **Bin 2 (mV)** | **Bin 3 (mV)** | **Bin 4 (mV)** | **Bin 5 (mV)** | **Bin 6 (mV)** |
| --- | --- | --- | --- | --- | --- | --- |
| **FDI** | 1.7 ± .1 | 1.79 ± .12 | 1.86 ± .12 | 1.88 ± .12 | 1.98 ± .12 | 2.14 ± .13 |
| **ADM** | 1.09 ± .09 | 1.14 ± .09 | 1.19 ± .1 | 1.2 ± .1 | 1.23 ± .1 | 1.31 ± .11 |

**Supplemental Table 2.** MEP amplitude raw data (mean ± standard error) from FDI and ADM muscles in the 6 bins in which we divided the 180 trials of the m-PAS.

**Supplemental Analysis 1**

To explore whether the CSE enhancement found during the m-PAS, reflecting a general increasing of M1 excitability, is persistent also after protocol’s administration, we run a ‘Trial type’ (static, movement) X ‘viewed Hand’ (left-hand, right-hand) X ‘Time’ (pre-PAS, post-PAS) X ‘Muscle’ (FDI, ADM) on MEP amplitude recorded during the action observation task and from which we derived our *motor resonance indexes*. Besides the significant quadruple interaction (*F*_1,80_ = 23.32, *p* < .001, *η_p_^2^* = .23), main factor ‘Time’ (*F*_1,80_ = .17, *p* = .685, *η_p_^2^* < .01), as well as interactions ‘Time X Muscle’ (*F*_1,80_ = 1.73, *p* = .192, *η_p_^2^* = .02), ‘Time X viewed Hand’ (*F*_1,80_ = 1.29, *p* = .259, *η_p_^2^* = .02), ‘Time X Trial type’ (*F*_1,80_ = 2.42, *p* = .124, *η_p_^2^* = .03), and ‘Time X Trial type X Muscle’ (*F*_1,80_ = 2.34, *p* = .13, *η_p_^2^* = .03) were not statistically significant. This pattern of results suggests that CSE modulation (and hence M1 excitability) were specific for the viewed hand, as already highlighted by the analyses reported in the main text. Crucially, CSE is not overall modulated before and after m-PAS administration.
